# Unexpected Properties of Short Genomic Tandem Repeats

**DOI:** 10.1101/165308

**Authors:** Irina Glotova, Michael Molla, Arthur L. Delcher, Simon Kasif

## Abstract

Length polymorphisms in genomic short tandem repeats have been implicated in a variety of diseases, most notably human neurodegenerative disorders. Expansions of tandem repeats are also associated with genomic instability in cancer. Our previous study of length-3 tandem repeats uncovered a surprising pattern in the length distribution of certain such repeats in the non-coding regions of the human reference genome: a bias towards repeats of length 3*n* - 1, (*n* > 3). That is, the observed frequency of repeats of this length in the human genome is higher than expected by chance based on the frequency of shorter repeats.

We have hypothesized that this pattern may be a general property of genomic DNA. If true, this could have implications with regard to the dynamics of repeat expansion generally. To test this hypothesis, we have analyzed the genomic sequences of a broad range of eukaryotic organisms as well as several complete human genomes and obtained a number of thought provoking results. We establish that this unexpected elevation in frequency of 3*n* - 1 long repeats is statistically significant. We also expanded this analysis to different classes of genomic regions and tandem repeats of length four and five. The specific pattern was found in 13 of the 20 organisms analyzed, including all chordate and insect genomes tested. The bias pattern, however, was not confined to a single branch of the evolutionary tree. For some genomes, such as *Drosophila melanogaster*, the repeat bias surprisingly was also identified in exons. The pattern is present in both small and large genomes. A similar pattern was also found in tetranucleotide and pentanucleotide repeats in the human genome. Another surprising property was identified for the flanking GC content for triplet repeats of length 3*n*. These findings indicate a puzzling new genomic phenomenon with possible evolutionary and disease-related implications.

## Findings

Short DNA sequences (1-6 bases long) repeated consecutively in a genome, are known as microsatellites or short tandem repeats (STRs). STRs have been reported to be linked with a variety of human neurological and neurodegenerative disorders [1-5], genomic instability in cancer [6] and are hypothesized to be associated with many other phenotypes. Nearly thirty diseases are associated with expansions in repeat sequences with a repeat unit of length 3 (3-STRs). These include Huntington’s disease, dentatorubral pallidoluysian atrophy, Machado-Joseph disease and numerous spinocerebellar atrophies [7-9]. Most of these diseases involve repeated glutamine codons (CAG). Tetra- and pentanucleotide repeats (CCTG, CAGG, ATTCT and AGAAT) are also associated with various neurological disorders. Typically, the lengths of these repeat regions vary throughout the population, with only those individuals having repeat lengths above a critical threshold level manifesting the disease.

STRs occur throughout the genome, both in coding and in non-coding regions. Microsatellite length distributions and the relative frequency of different repeat unit sequences have been shown to vary between species [2, 3]. STRs in different gene regions have been found to have various functional effects [10]. STR expansions and contractions in protein-coding regions can lead to a gain or loss of gene function via a frameshift mutation or an expanded toxic mRNA. STR variations in 5’-UTRs can affect gene expression by modifying transcription and translation mechanisms. STR expansions in 3’-UTRs may cause transcriptional slippage and produce expanded mRNA products, which can disrupt splicing and, potentially, other cellular functions. Intronic STRs can affect gene transcription, mRNA silencing, or export to cytoplasm. Triplet repeats located in UTRs or introns can also induce heterochromatin-mediated-like gene silencing. All these effects caused by STR expansions or contractions within genes can eventually lead to phenotypic changes and pathogenesis.

In a prior study [11], we observed a novel pattern in the length-3 repeat regions in the human reference genome [12] with possible functional implications for genomic instability in these regions. Specifically, tandem repeat sequences of length 3*n* - 1 (11 bp, 14 bp, 17 bp, etc.) are substantially more abundant than corresponding shorter length sequences (for example, 3*n* - 2 sequences: 10 bp, 13 bp, 16 bp, etc.). We called this the *saw-tooth* pattern because of the jagged pattern formed when the frequencies of repeat lengths are plotted as in Figure 1A. Here we provide a rigorous definition of this saw-tooth pattern and use this definition to ascertain which genomes from a broad range of organisms exhibit this genomic phenomenon. We also validate the presence of the saw-tooth pattern in additional sequenced human genomes.

**Figure 1.**
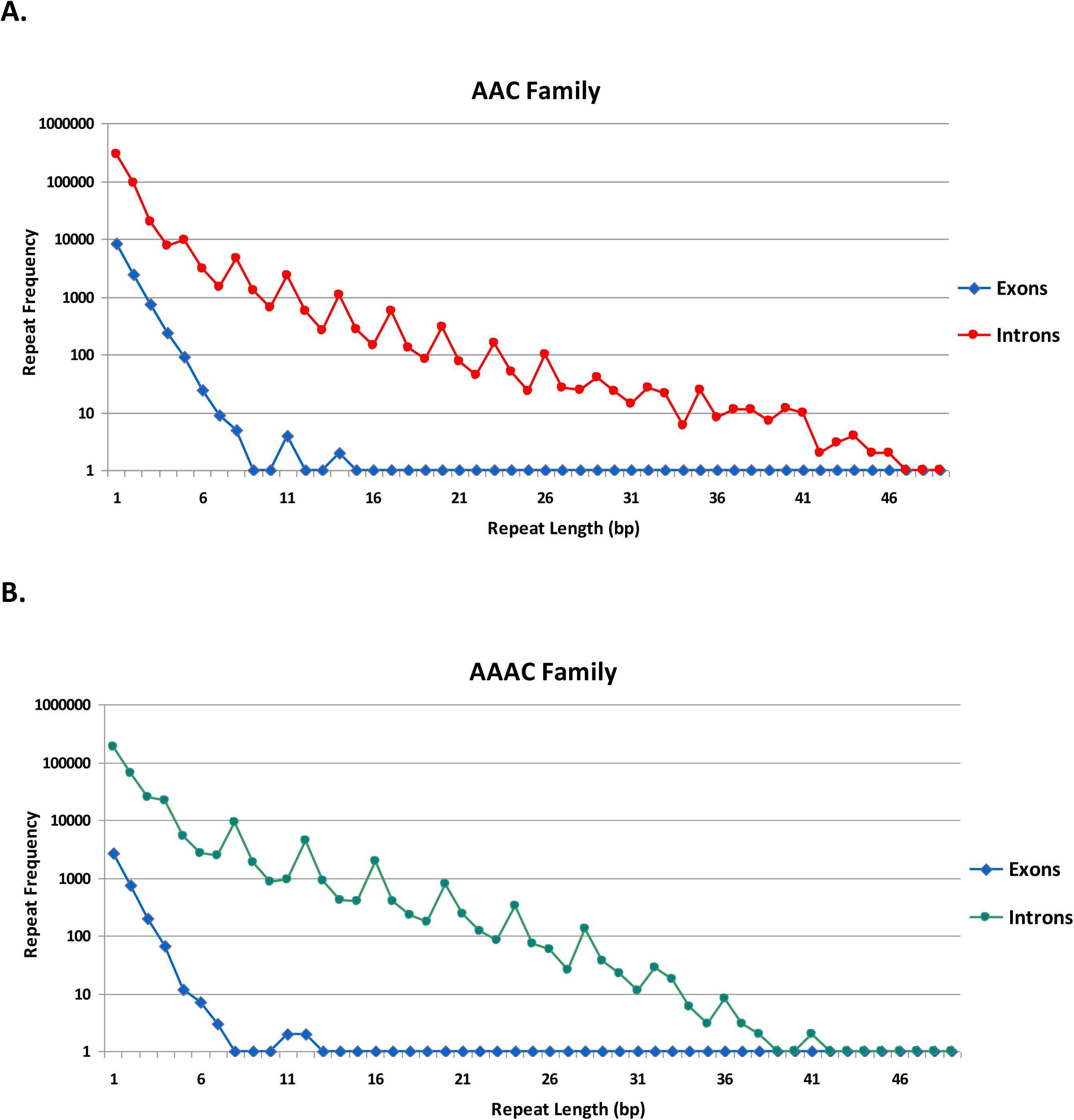
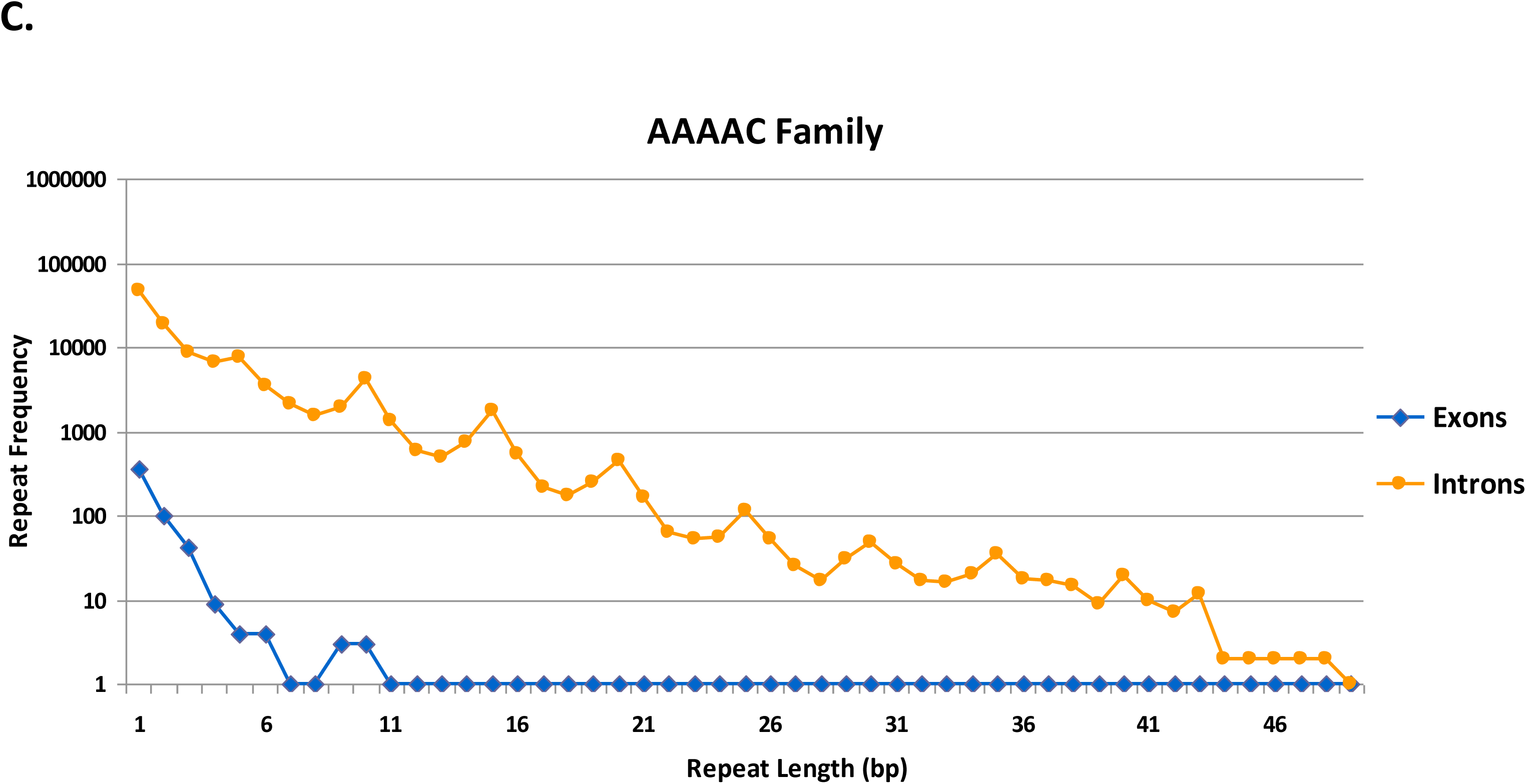
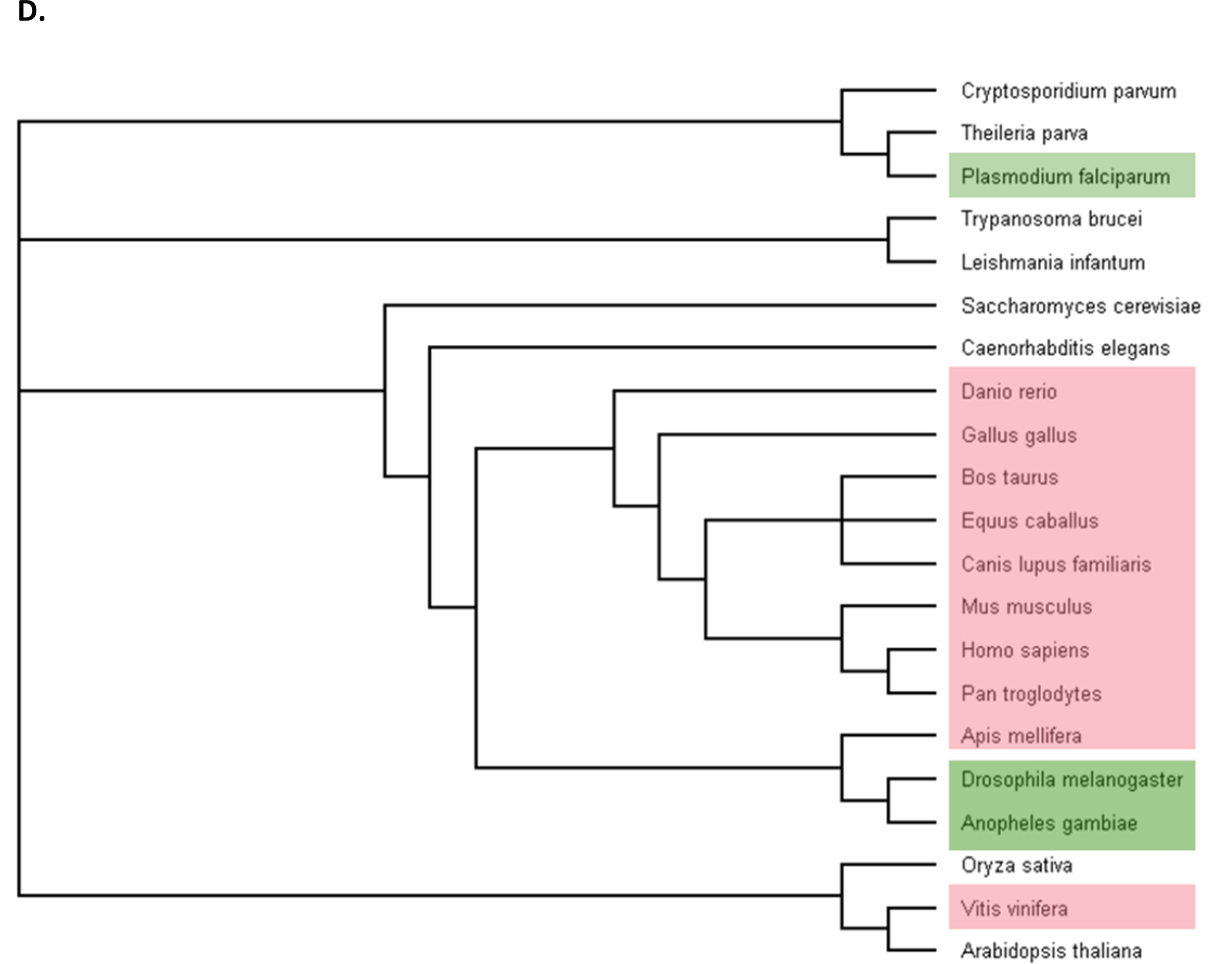
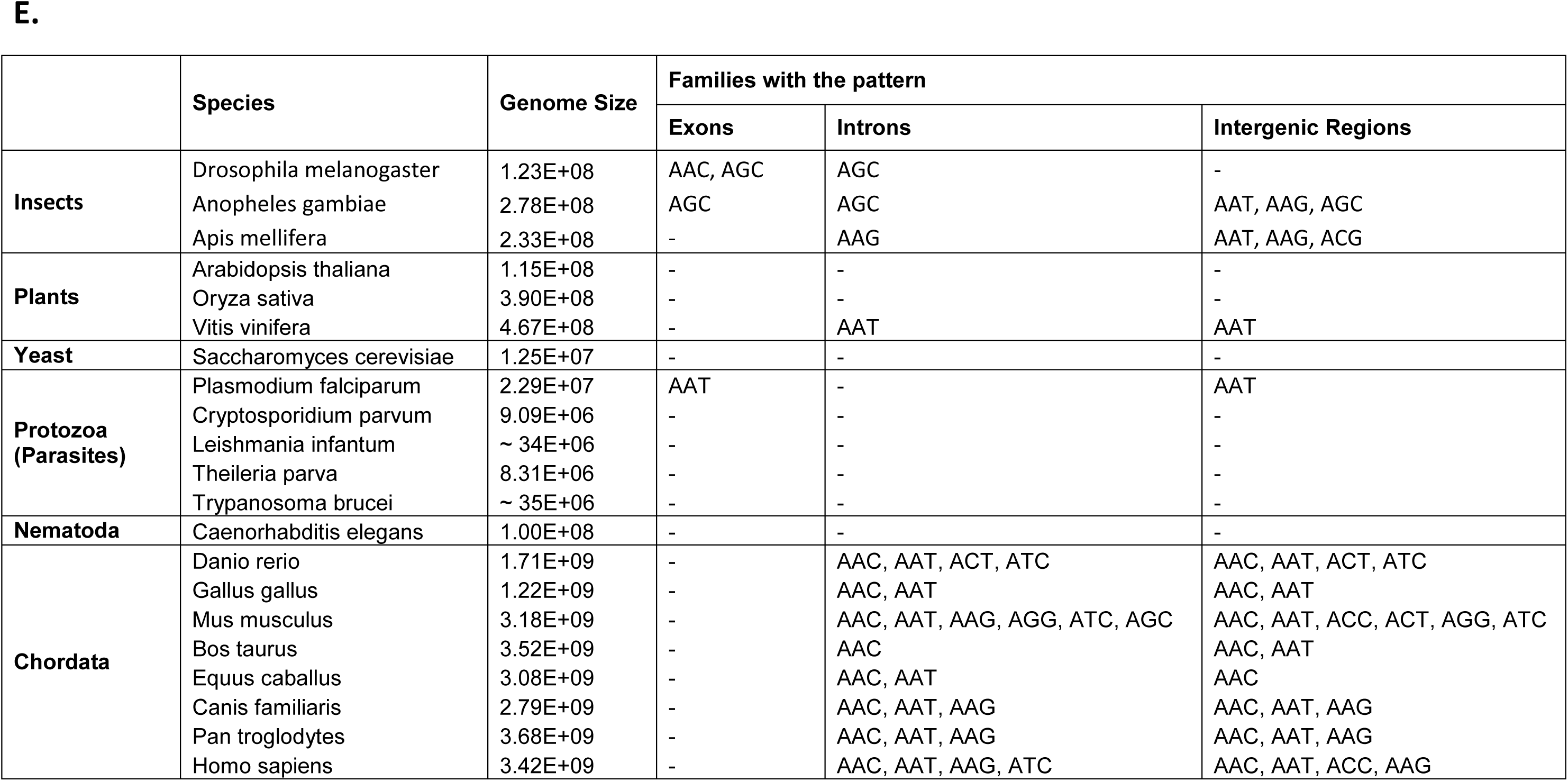
Repeat length bias in **(A)** tri-, **(B)** tetra- and **(C)** pentanucleotide repeats. Example of the saw-tooth pattern occurrence in AAC, AAAC and AAAAC repeat families (introns). **(D)** Taxonomic tree of the different genomes analyzed in the study. Genomes for which the saw-tooth pattern was identified in introns or/and intergenic regions are colored in pink, genomes for which the saw-tooth pattern was found in exons as well – in green. **(E)** List of the triplet repeat families in different genomes in which the saw-tooth pattern is present.

We analyzed the genomic composition and abundance of trinucleotide exact repeats (with minimum length of 7 nucleotides) in exons, introns and intergenic regions of the human reference genome (assembly GRCh37) [12], as well as the Korean [13] and Venter [14] genomes. We used Fisher’s exact test [15] for measuring whether the saw-tooth pattern exists in a particular family of repeats more than expected by chance. We define a family of repeats to be sequences whose repeat units are the same after either cyclic shift or reverse complementation. For example, the AGC family of triplet repeats includes all repeats in which the repeat unit is either AGC, GCA, CAG, GCT, TGC or CTG. For triplet repeat families, the test for the saw-tooth pattern is defined using the following procedure. Let *f(L)* be the number of occurrences of repeats of length *L*. For each type of repeat length, 3*n*, 3*n* - 1, 3*n* - 2 (*n* > 2) we measured the frequency of observed repeats of this length. Then for each length L, we computed the difference in the number of occurrences of repeat length L and L-1, *D(L) = f(L) - f(L-1)*. The differences were either positive or negative. We tested the hypothesis that the number of observed positive differences in all positions L = 3*n* - 1, for L > 7 is more than expected by chance using Fisher’s exact test (p-value < 0.05) (see Supplementary table).

In introns of the human reference genome, this definition showed the saw-tooth pattern to be present in AAC, ATC, AAT and AAG families (Supplementary figure 1), with AAC (Fig. 1A) being the most prominent and abundant one. In intergenic regions, the pattern could be seen in AAC, AAT, ACC and AAG families. The absence of the pattern in ACG and CCG families can be explained by the fact that very few repeats from this family were identified both in introns and intergenic regions. Additionally, CCG repeats, which exhibit considerable potential to form hairpins and quadruplexes, may also be selected against because they could affect the secondary structure of the pre-mRNA molecule, modulate the efficiency and accuracy of splicing, and interfere with the formation of mature mRNA [10]. No pattern was identified in the exons of the human reference genome. As expected, the analysis of the triplet repeats in the other two human genomes (Korean and Venter’s) showed that the saw-tooth pattern is present in the introns and intergenic regions of both genomes in the same set of repeat families. As these sequences were obtained using a variety of sequencing technologies, in addition to verifying that this pattern appears across the human population, this finding virtually eliminates the possibility that it is simply an error or a sequencing artifact. We also checked if some combinations of the repeat start and end are more prevalent for repeats of length 3*n* - 1 (see Supplementary figure 2). The number of repeats with combination 1:2 (for example, for AAC repeat family it starts and ends with an A) is considerably higher than the others. Therefore, it appears that this particular combination triggers the saw-tooth pattern.

**Figure 2.**
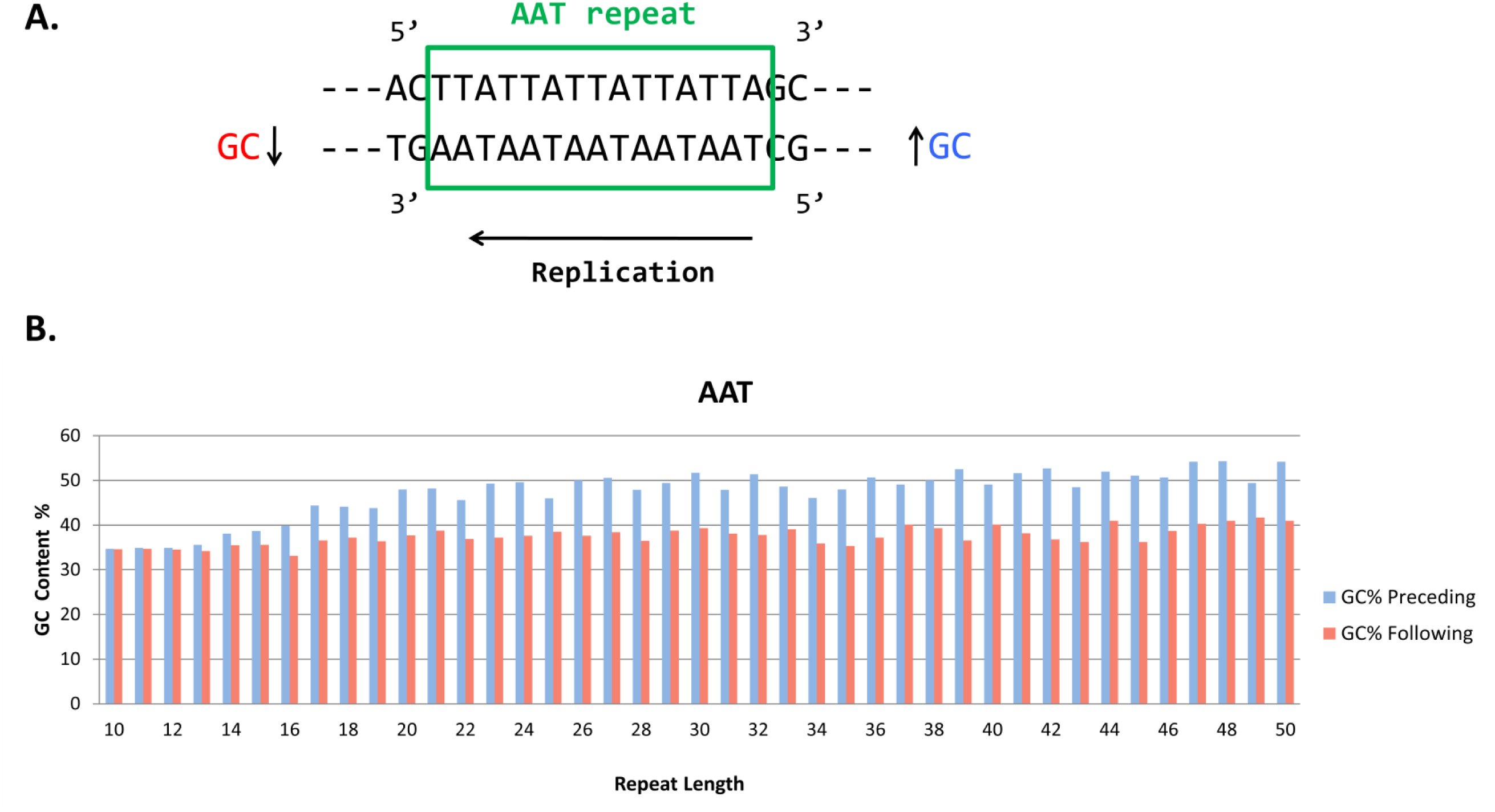
GC content bias for repeats of length 3*n*. **(A)** Explanatory figure. GC content preceding the repeat – blue, GC content following the repeat – red. The arrow shows the direction of replication. **(B)** GC content bias for AAT repeat family. GC content was calculated for 100 bp preceding/following the repeats.

Next, we sought to determine whether or not the saw-tooth pattern was limited to triplet repeats or if tetranucleotide and pentanucleotide repeats showed similar characteristics. We used the same method to analyze the genomic composition and abundance of tetra- and pentanucleotide exact repeats (with minimum length of 8 and 10 nucleotides, respectively). Analysis of the content of tetranucleotide repeats in the human reference genome showed that in introns a 4*n* - 1 saw-tooth pattern (analogous to the 3*n* - 1 pattern described above) was present in AAAC, AAAG, AAAT and AGAT families (4 out of 33 families). The most prominent of these, AAAC, is shown in Figure 1B. The rest of the families can be found in Supplementary figure 3. In intergenic regions the saw-tooth pattern was found in AAAC, AAAG, ACTC and AGAT families. Of the 102 pentanucleotide repeat families, only the three most abundant exhibit 5-periodic patterns. The 5*n* - 1 saw-tooth pattern was identified in the AAAAC repeat family (Figure 1C). For the other two most abundant families (AAAAG and AAAAT) more complex patterns are present (see Supplementary figure 4).

The presence of the saw-tooth pattern in human genomes led us to question how widespread the same pattern was in the genomes of other organisms and, in particular, if it was confined to certain regions in the evolutionary tree. To answer this question we analyzed the triplet repeat repertoire of 20 eukaryotic genomes which constitute different phylogenetic groups (plants, insects, yeast, Protozoa, Nematoda and Chordata). As shown in Figure 1D, the saw-tooth pattern was found to be present in introns and intergenic regions in all Chordata members analyzed. The highest number of repeat families with the pattern was found in *Mus musculus*. In plants the pattern was not found in *Arabidopsis thaliana* and *Oryza sativa* but was identified in both introns and intergenic regions of the *Vitis vinifera* genome. In the case of insect genomes the analyzed pattern is present in the introns of all three species and in the intergenic regions of *Anopheles gambiae* and *Apis mellifera*. Surprisingly, the saw-tooth pattern was also found in the exons of *D. melanogaster* and *A. gambiae* for the AGC repeat family. *D. melanogaster* protein-coding regions were shown to be enriched in repeats from AGC family [15] which partially explains this finding. No saw-tooth pattern is present in Nematoda (*Caenorhabditis elegans*) and yeast (*Saccharomyces cerevisiae*) genomes. Still, for Protozoa one out of five members analyzed (*Plasmodium falciparum*) has the pattern in both exons and intergenic regions (AAT repeat family). Similarly to *D. melanogaster* case, *P. falciparum* protein regions are known to have high AT content and to be specifically enriched in asparagine (AAT) repeats [16, 17]. Detailed findings are shown in Figure 1E.

Based on these results, we can conclude that the presence of the pattern seems not to depend directly on genome size. The pattern is present in both small and large genomes. Additionally, the occurrence of the saw-tooth pattern throughout the phylogenetic tree (Fig. 1D) suggests that this pattern does not have a single genetic origin. The cause or functional implications of this pattern are still not known.

We conjecture that this phenomenon could arise as the result of an in-phase repeat expansion mechanism such as DNA slippage [9]. If there were a minimum length, *m*, at which such an expansion could take place and, for at least some repeat families, this was a hard limit, *i.e*, there is an abrupt shift between repeats of length *m* – 1 and repeats of length *m*, the saw-tooth pattern would arise as an echo of the minimum length for expansion. That is, if some minimum length of repeat is required for a particular mode of repeat expansion, we would expect to see an abundance of repeat regions of length *m* + *nt*, where *m* is this minimum length for repeat expansion, *t* is the length of the repeat unit and *n* is any positive integer. Single-base expansions will still generate, in diminishing quantities, repeat lengths of *m* + 1, *m* + 2, *m* + 3, …. However, lengths of *m* + *t*, *m* + 2*t*, *m* + 3*t*, …, *m* + *nt* could occur more frequently given such a mechanism of repeat expansion. Our finding of a 3*n* - 1 pattern in some 3-STRs starting at length 11 suggests that, in the case of triplet repeats of certain repeat families, *m* = 8 bases might be the minimum length for this type of expansion.

Further study will be required to experimentally test our hypothesis that the saw-tooth pattern is caused by a length-thresholded mechanism of in-phase repeat expansion. The lack of a clear ancestral origin of this pattern suggests that there may be multiple such mechanisms. It may even be a common strategy for increasing genomic diversity throughout the living world. A better understanding of this mechanism would have implications in the understanding of a wide variety of serious genetic disorders associated with repeat expansion.

Another interesting property of genomic tandem repeats that was discovered in this study is related to the flanking GC regions. A correlation between the relative expandability of triplet repeats and the flanking GC content (presence of G and C nucleotides) has been observed by Brock *et al*. [18]. We mined the flanking GC content of triplet tandem repeats in the human genome to evaluate if there is any significant difference between the GC content before and after different classes of repeats (Fig. 2). Surprisingly, for AAT and AAC repeat families the GC content preceding the repeat is higher than the GC content following the repeat (Fig. 2B). This pattern is most prominent for repeats of length *3n* and is observed for flanking GC regions of various lengths (10, 20, 50, 200 bp). Integrating the replication direction with the increased difference in GC content in preceding and following flanking may stochastically affect the probability of repeat expansions. Thus the analysis of these patterns preceding and following a STR region and STR length can provide an insight into the replication direction that may allow us eventually to segment the human genome into replication regions.

## Authors’ contributions

All authors participated in the design of the experiments. IG performed all the experiments. AD and IG wrote all the software. All authors analyzed the results. IG wrote the paper and produced figures. All authors edited and approved the manuscript.

## Acknowledgements

The authors thank Charles Cantor, John Spouge and Kamila Naxerova for several useful discussions. SK was supported in part by grant NIH U54-LM008748.

